# SeqDesk: a sequencing-facility management system for standards-compliant and FAIR (meta)data submission

**DOI:** 10.64898/2026.08.05.743014

**Authors:** Philipp C. Münch, Gary Robertson, Alice Carolyn McHardy

## Abstract

Achieving FAIR compliance requires both standardized metadata and infrastructure for data deposition, yet in practice a large fraction of sequencing studies is still published without the persistent, standards-compliant metadata that reuse depends on. Collecting MIxS-compliant metadata is complex: environment-specific checklists can contain hundreds of fields, and the effort is magnified when metadata is assembled retrospectively at publication time rather than captured throughout the project. We developed SeqDesk, an open-source data management system for sequencing facilities that is designed so that FAIR-compliant public data is produced as the natural output of routine operations. Its current scope is microbial sequencing data, covering metagenomes as well as isolate genomes, for which it supports the corresponding MIxS checklists. SeqDesk gives a sequencing facility a configurable order-and-tracking system for sequencing projects, captures and validates MIxS-compliant metadata aligned with ENA checklists at project initiation, runs bioinformatics analyses through Nextflow pipelines, and brokers submission to the European Nucleotide Archive, all within the institution’s own infrastructure. By embedding standards-compliant metadata capture into the sequencing-facility workflow rather than bolting it on at submission, SeqDesk shortens the path from sample to reusable public data. The underlying checklist model is generic, so support can be extended to further data types and metadata standards beyond the microbial domain. SeqDesk is free and open source under the Apache 2.0 licence and available at https://seqdesk.org, with a live demonstration at https://seqdesk.org/#demo.

## Application note

The exponential growth of microbiome sequencing data has created unprecedented opportunities for understanding microbial ecosystems. With recent advances in AI, there is increasing demand for structured, high-quality metadata to enable automated data processing, cross-study comparison, and large-scale meta-analyses. The FAIR principles provide community guidelines to make data Findable, Accessible, Interoperable, and Reusable, and are now widely adopted in genomic data stewardship (Wilkinson et al. 2016). The MIxS (Minimum Information about any (x) Sequence) family of checklists developed by the Genomic Standards Consortium (GSC), including MIMARKS, MIGS, and MIMS, offers a robust basis for capturing environmental context, sampling protocols, and experimental designs (Bowers et al. 2017; Yilmaz et al. 2011). However, adoption remains challenging in practice. Best practices for data deposition often demand substantial effort with no immediate reward, and microbiome-specific tools for data sharing are still lacking, resulting in adoption gaps (Huttenhower, Finn, and McHardy 2023). The scale of this gap is easy to underestimate. We used the ENA Portal API to count all public sample records in the European Nucleotide Archive, grouped by the year in which they were first made public, using an exact whole-archive tally from a snapshot taken in late June 2026. In the most recent complete cohort, comprising roughly 4.95 million samples first made public in 2025, contextual metadata coverage was sparse: about 26% recorded geographic coordinates, 12% recorded a standardized MIxS environment field, and a collection date and country were each present in little more than half of the records. A reporting standard was near-universal in name only: 99% declared one, yet about a third used the permissive ‘Generic’ package, which mandates no geographic, environmental, or host context. Although the number of samples first made public each year has grown by roughly three orders of magnitude since 2008, contextual metadata coverage has improved unevenly and remains incomplete. Consequently, although the fraction of poorly contextualized records has not necessarily increased, their absolute number has grown substantially. Among the more than 40 million public samples first released between 2008 and 2025, many remain effectively ‘dark’, with sequence data lacking the metadata needed for reuse. Full sources and methods, including the exact ENA Portal queries and how missing-value placeholders are excluded, are given in the **Supplementary Material**. In addition, while community standards such as MIxS are widely recognized, their practical uptake remains inconsistent, leaving researchers uncertain about which fields are essential or how to apply controlled vocabularies effectively. These challenges are compounded by the fact that the perceived cost-benefit balance of detailed metadata annotation often discourages compliance, unless tools can shift these trade-offs by reducing effort (Sheffield, LeRoy, and Khoroshevskyi 2023). Furthermore, metadata collection frequently occurs during manuscript preparation to meet publication requirements. At that point, researchers may struggle to provide comprehensive documentation beyond the minimum requirements. This timing, which is often months or years after data generation, means that crucial experimental details may be incomplete or difficult to reconstruct. Collecting complete and well-structured metadata from the start is far more effective than attempting to improve poorly collected metadata retrospectively (Hughes et al. 2023).

Here, we introduce SeqDesk, an open-source web application designed for deployment within sequencing facilities so that institutions retain full control over infrastructure, data and access. At its core, SeqDesk is a sequencing-facility management system: it provides a standard, configurable order-and-tracking process for sequencing jobs, covering project intake, sample and library registration, run status and delivery. Administrators can adapt the workflow to their procedures by defining order-, study-, and sample-level fields and selecting the sequencing technologies, MIxS checklists, and optional modules offered to users. Unlike a general-purpose laboratory information management system (LIMS), SeqDesk is scoped specifically to the sequencing-facility workflow and to producing standards-compliant, submission-ready data. Built into that same workflow, it captures and validates MIxS-compliant metadata at project initiation through an intuitive interface with controlled-term suggestions and real-time checks, links this information to automated analysis via nf-core/Nextflow workflows (Ewels et al. 2020), and brokers submissions to ENA, so that standards-compliant, FAIR-ready public data is produced as a by-product of routine operation rather than a separate task. Researchers submit sequencing requests and metadata through the web front end, and sequencing facility staff use administrative tools to manage orders, update sample status, launch pipelines, and finalize public data deposition. SeqDesk is implemented in Node.js and TypeScript with a React interface, and is developed openly with continuous integration and an extensive automated test suite.

The platform provides a configurable sequencing-management system that covers the full operational workflow from project intake to data registration and upload. In the user frontend, researchers begin by creating an account and generating sequencing orders by specifying experimental parameters, sample information, library details, and, where configured, MIxS metadata. Existing samples can then be grouped into a study through the five-step workflow shown in **Fig. 1A**. During study creation, users select an appropriate ENA checklist derived from MIxS (**Fig. 1B**) and enter the corresponding environment-specific fields in the metadata table (**Fig. 1C**). The checklist definitions are represented as structured, versioned templates. Facility administrators can check the SeqDesk registry for updated ENA definitions, review added, removed, changed, and newly required fields, and apply an update. New studies record the active checklist version when they are created, and recent earlier definitions are retained so that existing studies can continue to use the version against which their metadata were entered. The templates are rendered in a spreadsheet-like interface with field validation, term suggestions, and context-aware help messages (**Fig. 1C**). For bulk sample entry, SeqDesk generates an Excel template from the configured per-sample fields. Users can complete or edit the template and upload it in the sequencing-order or study workflow. Before import, SeqDesk reports unmapped columns and row-level validation errors or warnings. On the sequencing facility side, an administrative interface mirrors this workflow and allows staff to process incoming orders, update and correct project or sample information, and modify metadata entries (**Fig. 1D**). Sequencing facility staff can also initiate processing pipelines from within the system. Facilities can tailor SeqDesk to their procedures without modifying the source code. Three form builders allow administrators to configure the sequencing-order questionnaire, the study questionnaire, and the per-sample fields recorded when samples are assigned to a sequencing run. Questions may be collected once per order or study, for every sample, or solely for internal sequencing facility use; their labels, help text, required status, choice lists, validation rules, visibility, and order are configurable. General-purpose field types can be combined with prebuilt domain-specific modules for sequencing technologies and barcodes, ENA-oriented organism taxonomy and sample identifiers, MIxS checklist selection and environment-specific metadata, and structured funding and billing information. Facilities can also configure which MIxS checklists, sequencing technologies, and optional modules are offered.

**Figure 1.**
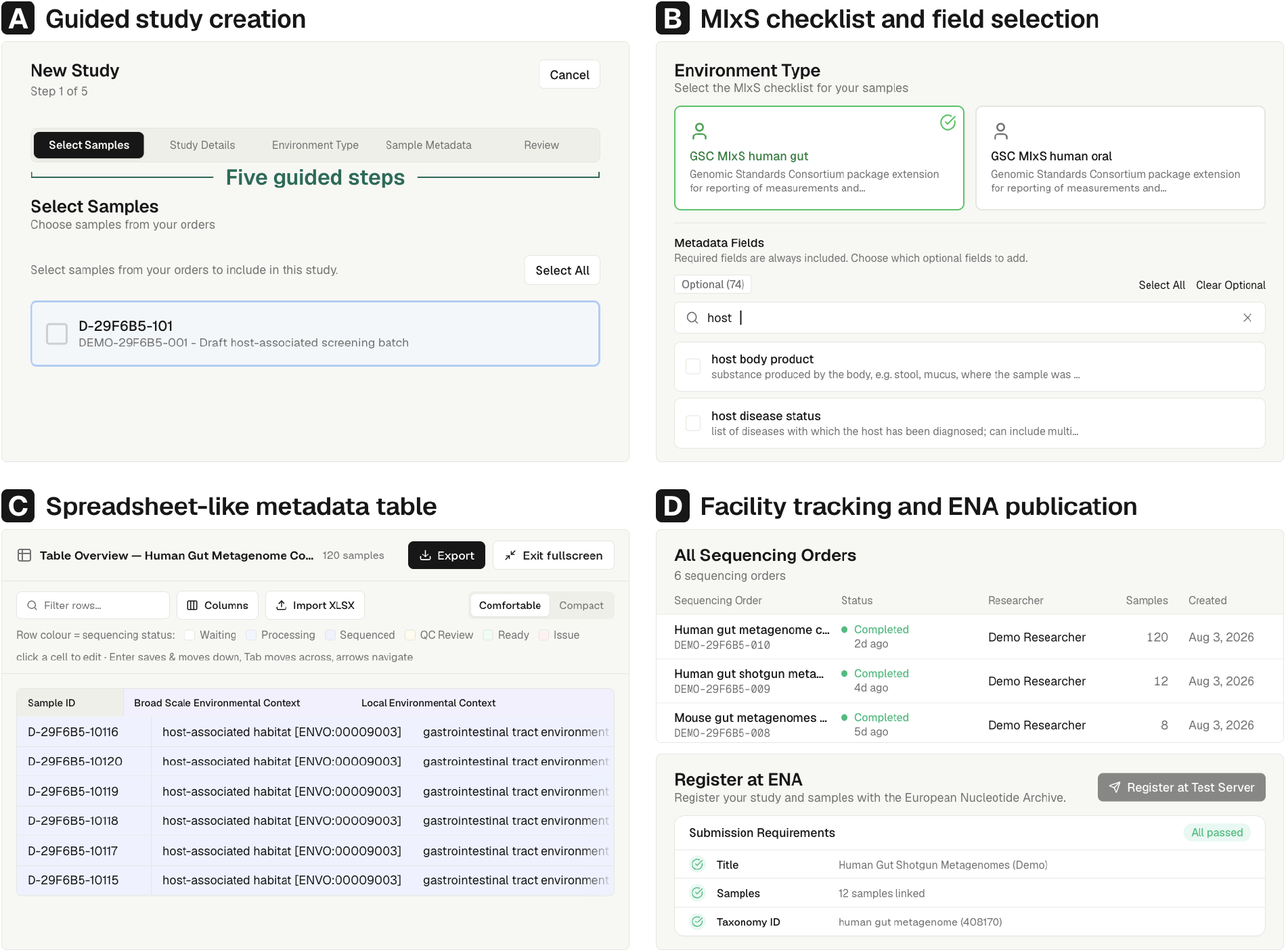
SeqDesk platform overview and key interfaces. (A) Study creation follows a five-step workflow comprising sample selection, study details, environment type, sample metadata, and review, allowing standards-compliant metadata to be captured while the study is defined rather than reconstructed at submission. (B) The registry-driven MIxS selector offers environment-specific checklists; once a checklist is chosen, required fields are fixed and the 74 optional fields in the checklist shown can be searched by name (here, host). (C) A spreadsheet-like metadata table supports direct cell editing with keyboard navigation and XLSX import and export; column tints mark field provenance, row tints mark sequencing status, and controlled-term suggestions are provided where applicable. (D) The facility workspace tracks sequencing orders by status, researcher, and sample count and supports ENA registration followed by data submission through subMG once all requirements are met.

In addition to managing the sequencing workflow itself, the platform allows facilities to execute downstream bioinformatic analysis on the sequencing data generated within each project. Because the entire system is deployed within the institution’s environment and can integrate directly with its existing compute infrastructure, it can use the same local resource to execute tasks such as assembly and binning of metagenomes. The platform connects to an on-site Nextflow instance with optional Slurm or HPC profiles, allowing staff to launch the nf-core/mag workflow directly from the interface, with all run provenance automatically linked back to the corresponding project and samples. Pipeline progress, logs, and QC reports are recorded and attached to each project, ensuring full traceability from raw reads to assembled metagenomes. Each pipeline is described by a manifest that declares its inputs, the sample sheet it expects, its configuration, and its outputs, and a single manifest-driven executor runs it either locally or on a Slurm cluster via Nextflow, so facilities can add any Nextflow pipeline (nf-core/mag being one ready-to-run example, alongside read cleaning and a lightweight test-data pipeline) and incorporate their preferred analysis standards at institutional scale.

SeqDesk also functions as a dedicated data-broker system for ENA submissions. The current broker targets ENA, reflecting the European research context in which SeqDesk was developed and the availability of established Webin and subMG interfaces. Direct submission to NCBI SRA is planned as a future extension. Sequencing facilities configure their ENA Webin credentials once in the administrative dashboard, after which studies and samples can be registered directly from the facility interface. Because sequencing orders, metadata, and pipeline outputs are all managed within the same system, the broker can link both raw reads and downstream analysis results with their corresponding samples during the submission, ensuring that ENA records capture all relevant data. Raw reads, metagenome assemblies, and optional genome bins can be submitted to ENA through the integrated subMG workflow, with returned accession numbers recorded on the corresponding SeqDesk records (Tubbesing, Schlüter, and Sczyrba 2025).

SeqDesk is designed to be simple to deploy and maintain. The core application installs natively with a single command (npm i -g seqdesk), a one-line installer, or from source, and requires only Node.js and PostgreSQL; it checks its prerequisites up front and, on any problem, stops with a specific, actionable message rather than failing silently. Running the analysis pipelines additionally requires a workflow runtime (Nextflow, with Conda or containers providing each pipeline’s tools); a scheduler such as Slurm is supported for cluster-scale execution but is not required, since pipelines also run locally. Access is controlled by locally-maintained user accounts; compute requirements for the portal are minimal, while those for pipeline processing are sample-dependent. The published software is continuously tested: over 3,600 automated tests run on every change, and continuous integration installs the release on a clean machine and verifies that it starts, applies its database migrations, and supports login, alongside versioned releases and a plain-language changelog. User and operator documentation is maintained at https://seqdesk.org/docs and covers installation, configuration, sequencing orders and studies, sequencing files, pipelines, ENA submission, administration, updates, and troubleshooting. Within the application, a Help & Guide page explains the main workflow, user roles, and the location of common tasks, while order and study forms provide field-specific help and validation guidance. A demonstration environment that requires no installation is available at https://seqdesk.org/#demo. Source code and tagged releases are available under the Apache License 2.0 at https://github.com/hzi-bifo/SeqDesk, with versioned releases archived on Zenodo at https://doi.org/10.5281/zenodo.21806366.

## Discussion and Conclusions

Integrating metadata collection into the sequencing-facility workflow addresses the gap between sequencing operations and public data deposition. SeqDesk captures metadata during order intake and sample or library registration, before sequencing begins, rather than requiring researchers to reconstruct it after sequencing or during manuscript preparation. Metadata can therefore be checked against the selected ENA checklist while the study is active and before public submission. By connecting researchers, sequencing operations, bioinformatic workflows, and public archives through the sequencing facility, SeqDesk makes standards-aligned metadata capture part of routine data generation.

There are several platforms that support metadata collection and, in many cases, data submission. Collaborative OPen Omics (COPO) brokers metadata annotation using community standards and submits datasets to appropriate public repositories while tracking accessions (Shaw et al. 2020). METAGENOTE provides MIxS-guided metadata capture and automates submission to NCBI BioProject, BioSample, and SRA (Quiñones et al. 2020). Qiita, primarily a platform for large-scale meta-analysis and data reuse, also supports uploading study/sample/preparation metadata and raw sequence files and can broker public deposition to ENA through its “Make data public” workflow (Gonzalez et al. 2018). DataHarmonizer is a browser-based template and validator that enables structured entry, checks and transformation to submission-ready formats (Gill et al. 2023). SeqDesk differs from these systems by integrating MIxS-compliant metadata capture with sequencing-facility order tracking, downstream analysis, and ENA submission in a single locally deployed platform. This integration enables metadata collection near project initiation while keeping it linked to facility operations. In the broader environmental and biodiversity domain, platforms such as PANGAEA, the GFBio portal (Diepenbroek et al. 2014) and BEXIS2 (Chamanara et al. 2021) provide infrastructures for managing and publishing heterogeneous research metadata, but to our knowledge, they are not integrated into sequencing-facility workflows. Tools like the NMDC Field Notes app (Kalita, Cavanna, and Xu 2024) further highlight the importance of capturing structured, location-aware metadata directly at the time of sampling, typically guided by domain-specific schemata. SeqDesk occupies a complementary niche by embedding the capture and validation of MIxS-based metadata aligned with ENA checklists within sequencing-facility operations and linking this information directly to downstream Nextflow workflows, including nf-core and other workflows, and automated ENA submissions. At present, SeqDesk focuses on microbial metagenomes and genomes; because checklists are configurable rather than hard-coded, we intend to extend SeqDesk to further data types and metadata standards in future releases.

## Acknowledgements

We gratefully acknowledge the valuable input of the GSC Scientific Advisory Board. This work was supported by NFDI4Microbiota, funded by the Deutsche Forschungsgemeinschaft (DFG, German Research Foundation) under project number 460129525.

## Notes

### Competing Interest Statement

The authors have declared no competing interest.

