## Supplementary Material for "SeqDesk: a sequencing-facility management system for standards-compliant and FAIR (meta)data submission"

### S1. FAIR-gap metadata analyses: how the background numbers were generated

The figures cited in the introduction come from three complementary, archive-wide analyses of public sample records in the European Nucleotide Archive (ENA). The analyses use server-side aggregate counts returned by the ENA Portal API count endpoint (https://www.ebi.ac.uk/ena/portal/api/count) with result=sample. Only aggregate counts and percentages are published; no sample identifiers are retrieved, stored, or released. All public ENA sample records with a first_public date from 1 January 2008 onward are grouped by the calendar year in which they were first made public, using first_public>=Y-01-01 AND first_public<=Y-12-31. These are therefore first-public-year cohorts, not submission-year cohorts. For each field, the cohort total and populated-field count were queried and, where applicable, separate counts matching controlled missing-value terms were subtracted. The calculations therefore include every record matching the defined queries rather than a downloaded subsample, avoiding the project-clustering bias that could result from downloading the first N records of a year. Headline figures use the most recent complete first-public-year cohort, comprising approximately 4.95 million records in 2025; the partial current year is excluded. The values cited in the manuscript come from late-June 2026 retrievals (metadata completeness, 26 to 29 June; reporting standard, 25 June). Together, the 2008 to 2025 cohorts contained approximately 43.77 million public sample records.

#### Metadata completeness

For each cohort, we count records in which selected contextual fields contain a non-missing value. For text fields, this means that the field is present and does not match one of the INSDC controlled missing-value terms, including missing and values beginning with missing:, not collected, not provided, not applicable, or restricted access. This operational definition measures non-missingness, not the semantic correctness, accuracy, or ontology validity of a value. Geographic coordinates are counted through the latlon-typed location column; collection dates through a date-range filter on collection_date from 1900-01-01 to the retrieval date, meaning that earlier and future dates are excluded; the MIxS environment through the environment_biome column, after subtracting missing-value terms; and country through a populated country value after subtracting missing-value terms. In the 2025 cohort, approximately 26% of records contained coordinates and 12% contained a non-missing environment value; the corresponding proportions for collection date and country were approximately 56% and 59%, respectively.

#### Reporting standard

For each cohort, we examine the ncbi_reporting_standard column, which records the NCBI BioSample reporting package associated with a record. “Any standard” denotes a populated value, Generic is counted separately, and “non-Generic” is calculated as the populated count minus the Generic count; matching is case- and spelling-sensitive. In the 2025 cohort, approximately 99% of records had a populated reporting-standard value, about 33% of all records were labelled Generic, and about 66% had a non-Generic value. This analysis describes reporting-package declarations only. It does not imply that every non-Generic package is a GSC MIxS package or that the metadata requirements of the declared package were met.

**Volume versus context.**

For each first-public year, we compare the cohort size with the share carrying a non-missing MIxS environment value. Annual cohort size increased by roughly three orders of magnitude, from a few thousand records per year in the late 2000s to millions per year in 2025. Environment-field coverage rose from below 1% in the earliest cohorts to approximately 12% in 2014, then fluctuated between approximately 8% and 14% through 2025. Consequently, the absolute number, but not necessarily the fraction, of records lacking this field increased substantially. Because the field applies mainly to environmental samples and the denominator includes all sample types, this metric is influenced by archive composition and cannot by itself classify records as dark or unusable. It is one indicator of contextual-metadata availability.

Caveats and reproducibility. Each annual point includes all public ENA sample records matching the defined first_public date range during the stated retrieval period. Large projects and changes in archive composition can shift individual yearly values, and field applicability varies among sample types. The counts are server-side aggregates rather than estimates obtained through subsampling. The manuscript values refer to the late-June 2026 retrievals specified above. Live aggregate results and interactive charts are available at https://seqdesk.org/data under the metadata-completeness, reporting-standard, and deluge-vs-dark analyses. These live analyses are scheduled for weekly refresh and may therefore display newer values.
